# Targeting of CXCR4 by the naturally occurring CXCR4 antagonist EPI-X 4 in Waldenström’s Macroglobulinemia

**DOI:** 10.1101/2020.10.19.344812

**Authors:** Lisa M. Kaiser, Mirja Harms, Daniel Sauter, Vijay PS Rawat, Mirco Glitscher, Eberhard Hildt, Daniel Tews, Zachary Hunter, Jan Münch, Christian Buske

## Abstract

CXCR4 expression and downstream signaling have been identified as key factors in malignant hematopoiesis. Thus, up to 40% of all patients with Waldenström’s Macroglobulinemia (WM) carry an activating mutation of CXCR4 that leads to a more aggressive clinical course and inferior outcome upon treatment with the Bruton’s tyrosine kinase inhibitor ibrutinib. Nevertheless, little is known about physiological mechanisms counteracting CXCR4 signaling in hematopoietic neoplasms. Recently, the endogenous human peptide EPI-X4 was identified as a natural CXCR4 antagonist that effectively blocks CXCL12-mediated receptor internalization and suppresses the migration and invasion of cancer cells towards a CXCL12 gradient. Here, we demonstrate that EPI-X4 efficiently impairs growth of WM cells *in vitro* and *in vivo* and blocks their migration towards CXCL12. The CXCR4 inhibitory activity of EPI-X4 is accompanied by decreased expression of genes involved in MAPK signaling and energy metabolism. Notably, the anti-WM activity of EPI-X4 could be further augmented by the rational design of EPI-X4 derivatives showing higher binding affinity to CXCR4. In summary, these data demonstrate that a naturally occurring anti-CXCR4 peptide is able to interfere with WM growth, and that optimized derivatives of EPI-X4 may represent a promising approach in suppressing growth promoting CXCR4 signaling in WM.

## Introduction

The CXCL12/CXCR4 axis represents one of the most fundamental pathways regulating cell anchorage in the bone marrow (BM) and trafficking of cells to distant organs. This regulatory principle is highly conserved between species and also maintained in neoplasms, shaping the behavior of malignant cells. For example, CXCR4 is overexpressed in 25–30% of patients with acute myeloid leukemia (AML) and is associated with poor prognosis [1]. Functional assays have proven that the CXCL12/CXCR4 crosstalk is critical in regulating the interaction between normal hematopoietic stem cells (HSCs) but also leukemic stem cells (LSCs) and their niche [2]. Beside this, CXCR4 is vital for normal lymphocyte homing and trafficking and is closely linked to the pathobiology of several lymphomas such as chronic lymphocytic leukemia (CLL), diffuse large B cell lymphoma (DLBCL), follicular lymphoma (FL), marginal zone lymphoma (MZL), hairy cell leukemia (HCL) and mantle cell lymphoma (MCL) [1,3–8]. For example, high expression of CXCR4 is a hallmark of CLL cells compared to normal B cells [6,9] and is associated with advanced Rai stages [10]. Furthermore, a genome-wide screening of familial CLL revealed germline mutations in the coding regions of CXCR4 [11]. However, Waldenström’s Macroglobulinemia (WM) is probably the lymphoma subtype that is most closely linked to CXCR4 and its downstream signaling [12]. WM is an incurable B-cell neoplasm characterized by serum monoclonal immunoglobulin M (IgM) and clonal lymphoplasmacytic cells infiltrating the bone marrow. Recent years have succeeded to describe the molecular landscape of WM in detail, highlighting two recurrently mutated genes, *MYD88* and *CXCR4*; *MYD88* with an almost constant and recurrent point mutation (L265P) present in over 90% of patients and *CXCR4* with over 40 different mutations in the coding region, affecting up to 40% of patients, among them the most frequent mutation C1013G/A mutation, predicting a stop codon in place of a serine at amino acid position 338 (S338X) [13]. Intriguingly, both mutations are activating mutations leading to an indelible activation and perpetual signaling of the chemokine receptor in the case of CXCR4 [14–16]. These observations have shed light on the essential role of CXCR4 in WM and have paved the way for predicting treatment response to the Bruton tyrosine kinase (BTK) inhibitor ibrutinib and novel therapeutic approaches, which might be transferable to other CXCR4-driven diseases [17,18].

Our group has recently identified a naturally occurring CXCR4 inhibitor termed Endogenous Peptide Inhibitor of CXCR4 (EPI-X4). EPI-X4 is a 16-mer peptide that is produced at low pH via proteolytic cleavage of serum albumin by Cathepsin D and E. In contrast to the clinically approved CXCR4 inhibitor AMD3100/Plerixafor, EPI-X4 does not bind CXCR7 and has no mitochondrial cytotoxicity, suggesting lower side effects. Intense research has already shown that EPI-X4 is able to interfere with the crosstalk of CXCR4 and CXCL12, inhibits infection with CXCR4-tropic HIV-1 strains, mobilizes hematopoietic stem cells and is able to suppress migration of leukemic cells *in vitro* [19].

We now demonstrate that EPI-X4 interferes with CXCR4 signaling in WM, where it impairs malignant growth *in vitro* and *in vivo*. These data identify an endogenous protective mechanism that dampen the prosurvival signals of CXCR4 in WM, a key oncogenic driver in this disease.

## Results

### CXCR4 expression in patients with WM

First, we analyzed the transcription levels of *CXCR4* in primary WM patient samples according to their mutational status of *MYD88* and *CXCR4* based on a previously reported series of patients analyzed by RNA-Seq [20]. WM patients harboring the MYD88 L265P mutation expressed on average higher levels of CXCR4 compared to normal peripheral B cells and memory B cells (Fig. 1A). In contrast, CXCR4 expression in MYD88 wildtype (MYD88^WT^) WM patients was significantly lower compared to *MYD88* mutated (MYD88^Mut^) cases and normal circulating B cells. Finally, CXCR4 expression of patients carrying mutations in both *MYD88* and *CXCR4* (MYD88^Mut^/CXCR4^Mut^) was on average lower than that of cases carrying only a MYD88 mutation, but higher than MYD88^WT^/CXCR4^WT^ patients (Fig. 1A). We validated these findings using a second data set, comprising RNA-Seq analyses of 16 patients, all being MYD88 mutated with seven of these patients carrying a concurrent CXCR4 mutation [21]: as shown in Fig. 1B, CXCR4 was highly expressed both in CXCR4 mutated and non-mutated cases. Of note, there was a significant higher expression of HIF-1a and MAPK1 in CXCR4 mutated cases, associated with a clinical more aggressive course [22]. Taken together, these data demonstrate high expression of CXCR4 in WM cells harboring the MYD88 L265P cases independent of the CXCR4 mutational status and show high expression of genes associated with CXCR4 signaling and tumor growth particularly in CXCR4^Mut^ cases.

**Fig. 1:**
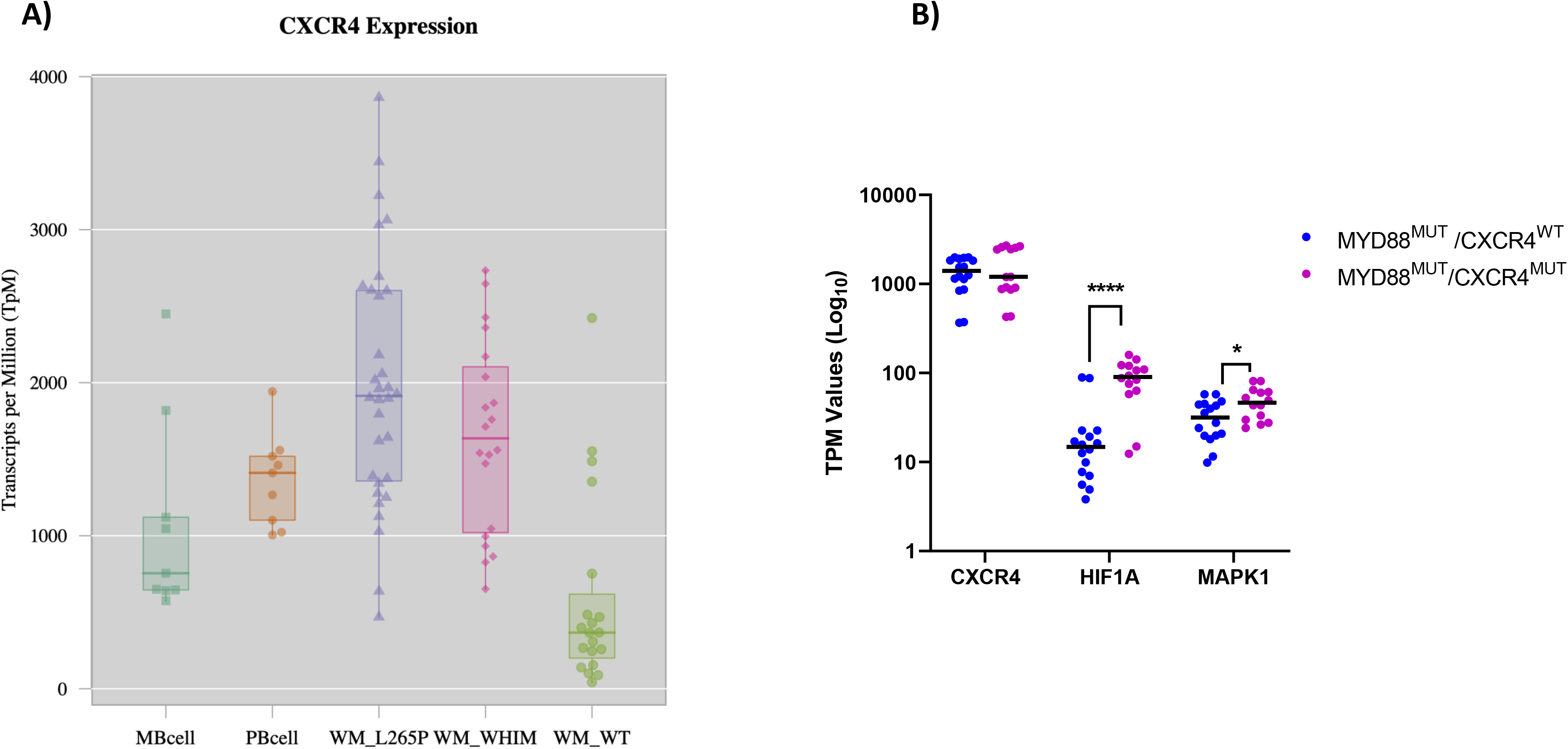
Expression of CXCR4 in WM patients. **(A)** Expression of *CXCR4* determined by RNA-Seq based on a data set of 57 WM patients compared to healthy B-cells [20]. MB cell = memory B cells; PB cell = peripheral B cells; WM_L265P= MYD88 mutated cases; WM_WHIM= MYD88 and CXCR4 mutated cases; WT = cases lacking MYD88 and CXCR4 mutations **(B)** Expression of CXCR4, HIF-1a and MAPK1 determined by RNA-Seq in 15 MYD88 mutated WM patients with (WHIM) and without (WT) CXCR4 mutation. * = p< 0,02; *** = p< 0,00002.

### The CXCR4 mutation S338X promotes cell proliferation in WM

As we had seen high expression of CXCR4 in both CXCR4 mutated and non-mutated WM patients, first we tested the impact of wild type and mutated CXCR4 on the growth of the WM cell line BCWM.1, which is MYD88^Mut^/CXCR4^WT^. To this end, BCWM.1 cells were lentivirally engineered to express wildtype CXCR4 (iso1 WT) or patient-derived variants carrying the mutations S338X, R334X, S339fs/342X or S339fs/365X. As shown in Suppl. Fig. 1, S338X showed a strong effect on clonogenicity with a 30% increase compared to the empty vector control. In contrast, the CXCR4 mutations R334X, S339fs/342X and S339fs/365X as well as the iso1 WT did not increase colony number as efficiently as CXCR4 S338X or even reduced colony numbers in the case of S339fs. CXCR4 S338X also significantly enhanced proliferation of this cell line *in vitro* compared to cells transduced with the vector control or the *CXCR4* WT gene (Fig. 2A). Finally, the growth promoting effect of the S338X mutation was confirmed after lentivirally engineered expression in CXCR4 wildtype MCWL-1 WM cells where constitutive expression of the S338X mutant induced a 3.5-fold increase in clonogenic growth compared to the empty vector control (Fig. 2B). Together, these data demonstrate that expression of the activating CXCR4 mutation S338X promotes cell growth of WM cells.

**Fig. 2:**
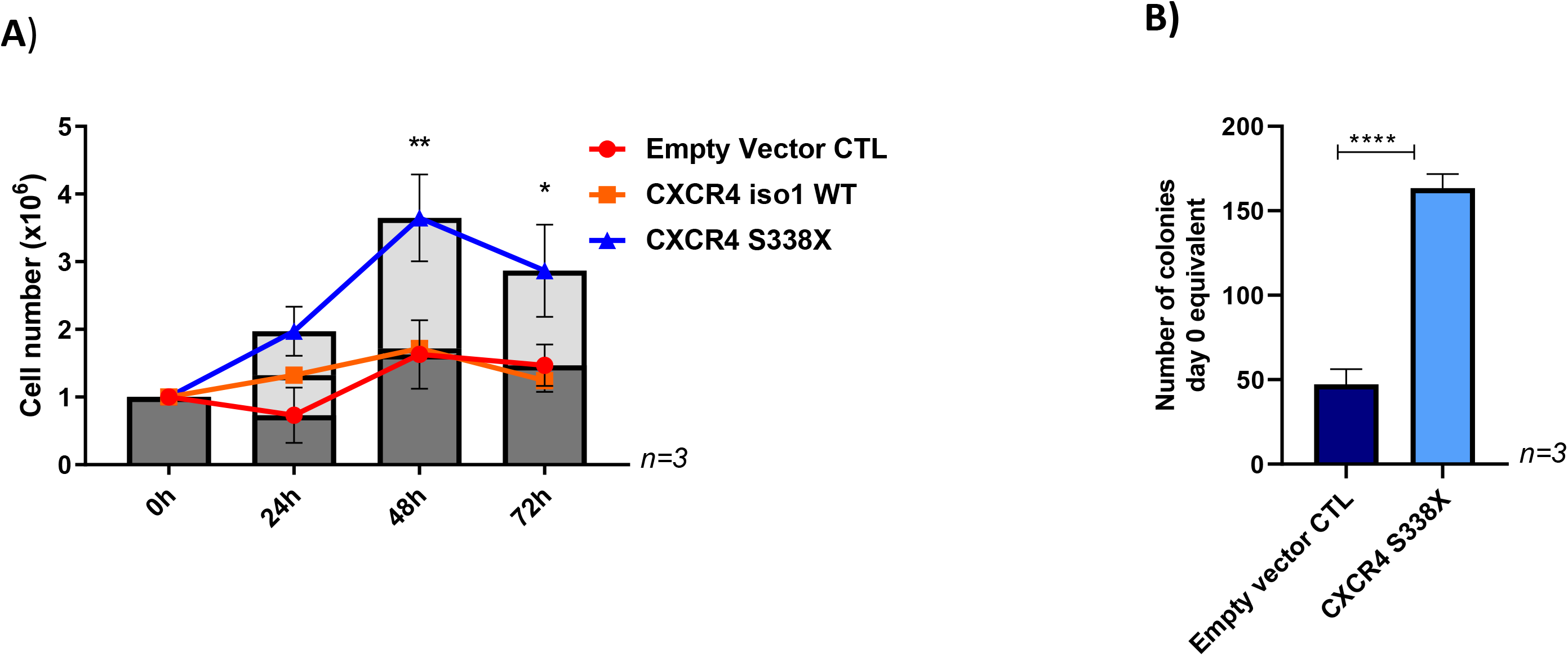
Impact of the CXCR4 S338X WM mutation on growth. **(A)** Increased proliferation of BCWM.1 cells transduced with CXCR4 S338X. Absolute cell numbers were determined after 24, 48 and 72 h using Trypan blue for live-dead staining. Mean values (± SEM) of three independent experiments are shown (** p<0.002; * p<0.03). **(B)** Increased colony formation ability of MCWL-1 cells transduced with CXCR4 S338X. Mean values (± SEM) of three independent experiments performed in duplicates are shown (*** p<0.0001).

### The natural CXCR4 antagonist EPI-X4 and its optimized derivatives impair growth of WM cell in vitro and in vivo

To investigate a potential role of the natural CXCR4 antagonist EPI-X4 in WM, we first tested its ability to bind and block CXCR4 variants observed in WM patients. Efficient and dose-dependent blocking of CXCR4 by EPI-X4 was previously documented in competition experiments using two different clones of CXCR4 antibodies that recognize different epitopes in the extracellular part of CXCR4: while the antibody clone12G5 binds to the epitope overlapping with the EPI-X4 binding site, 1D9 binds to an epitope in the CXCR4 N-terminus, which is not masked by the peptide. In line with this, efficient blockage of the 12G5 epitope by EPI-X4 was achieved at a concentration of 200 μM for all CXCR4 mutants tested (Fig. 3A). In contrast, the 1D9 epitope could not be blocked by EPI-X4 (Fig. 3B). These results were confirmed in MCWL-1 cells (data not shown).

**Fig. 3:**
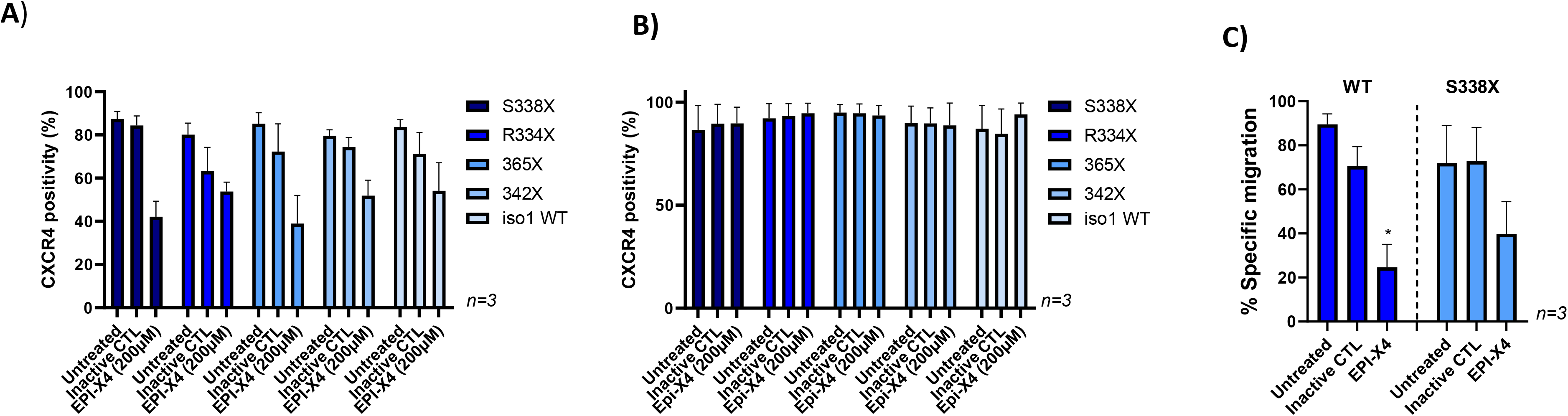
EPI-X4 blocks the CXCR4 12G5 epitope on BCWM mutant cells and impairs migration. BCWM.1 cells were preincubated with 200 μM EPI-X4 or inactive control peptide and then stained with the anti-CXCR4 antibody clones 12G5 **(A)** or 1D9 **(B)**. While EPI-X4 blocks binding of the anti-CXCR4 clone 12G5 to all CXCR4 variants tested, it did not affect binding of the 1D9 clone. Mean values (± SEM) of one exemplary experiment performed in triplicates are shown. **(C)** CXCL12-directed transwell migration of BCWM.1 cells expressing wildtype CXCR4 (WT) or CXCR4 S338X was monitored in the presence or absence of EPI-X4 (200 μM). Mean values (± SEM) of three independent experiments performed in duplicates are shown (*p<0.03 and ns, respectively).

Since EPI-X4 efficiently binds to CXCR4 mutants, we initiated functional assays testing the effect of this naturally occurring CXCR4 antagonist on migration of WM cells towards a CXCL12 gradient *in vitro*. Transwell assays revealed that EPI-X4 reduces migration compared to an inactive control peptide in BCWM.1 cells expressing wildtype or S338X CXCR4 by 65% and 45%, respectively (Fig. 3C).

To investigate whether rational design can further enhance the anti-CXCR4 and therefore anti-WM activities of EPI-X4, we took advantage of two EPI-X4 derivatives, termed WSC02 and JM#21. These peptides were generated from EPI-X4 by structure-activity-relationship (SAR) studies and subsequent synthetic modulations inserting mutations in the lysine residues [23] and were also able to efficiently block the epitope 12G5 at low concentrations (Suppl. Fig. 2). To test whether WSC02 is functionally active and superior to the parental EPI-X4 peptide, migration tests were performed using BCWM.1 wildtype cells. As shown in Fig. 4A, WSC02 was more efficient in blocking migration than EPI-X4. Analysis of lentivirally engineered BCWM.1 cells revealed that WSC02 also efficiently blocks migration of CXCR4 S338X expressing cells, albeit slightly less efficiently than cells transduced with wildtype CXCR4 (81% vs. 64%, respectively). JM#21 showed an even more pronounced activity, reducing migration of CXCR4 iso1 wildtype and CXCR4 S338X expressing WM cells by 100% and 93%, respectively (Fig.4B). To test the ability of EPI-X4 and WSC02 to impair lymphoma progression *in vivo*, MCWL-1 cells were incubated with one of the two peptides or the inactive peptide control *in vitro* and transplanted into NSG mice. While WSC02 significantly delayed engraftment and nearly doubled the median overall survival, EPI-X4 had no major impact (Fig. 4C). Taken together, EPI-X4 and its optimized derivatives are able to impair growth of WM cells expressing wild-type or mutated CXCR4 and the EPI-X4 derivative WSC02 even increased survival in a murine WM model.

**Fig. 4:**
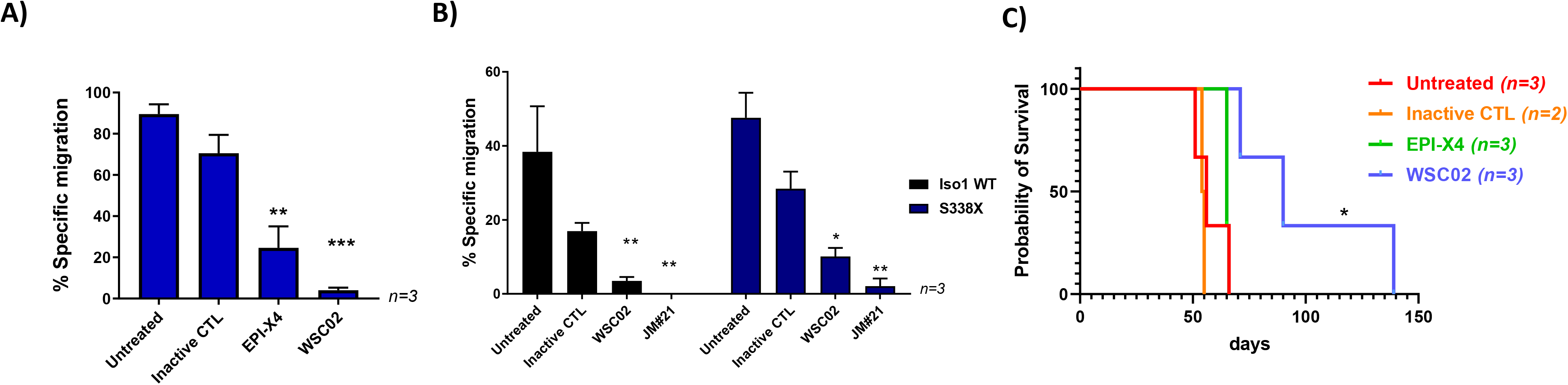
Optimized EPI-X4 derivatives impair migration and prolong survival of NSG mice transplanted with WM cells. **(A)** WSC02 reduces CXCL12-directed transwell migration of BCWM.1 WT cells at a concentration of 10 μM more efficiently than EPI-X4 at a concentration of 200 μM (Ordinary one-way Anova **p<0,06; ***p<0,0005). **(B)** WSC02 and JM#21 suppress migration of BCWM cells stably expressing CXCR4 iso1 WT or CXCR4 S338X at a concentration of 10 μM. All values represent mean values ± SEM of migrated cells relative to CXCL12-only treated cells from three independent experiments (*p=0,02; **p=0,006).**(C)** Survival of mice injected with MCWL-1 cells treated with the indicated peptides. Median survival: Untreated: 56 days; Inactive CTL: 54.5 days; EPI-X4: 65 days, WSC02: 90 days. Log-rank (Mantel-Cox) test: Untreated/Inactive control (n=5) compared to WSC02 (n=3) (*p<0,01)

### Inhibition of CXCR4 by EPI-X4 changes downstream signaling and gene expression in WM cells

To understand the mechanisms underlying the inhibitory effect of EPI-X4 and its optimized derivatives, we next sought to determine whether EPI-X4 and the optimized EPI-X4 derivatives have the ability to suppress ERK signaling in WM cells harboring different CXCR4 mutations, functioning as inverse agonists, independently of CXCL12. To this end, we monitored ERK phosphorylation at Tyr204 (ERK1) and Tyr187 (ERK2) in stably CXCR4 transduced BCWM.1 cells after CXCL12 stimulation using flow cytometry. EPI-X4 had a weak inhibitory effect in CXCR4 WT expressing cells with a more pronounced effect in CXCR4 S339fs mutated cells and complete resistance in cells carrying the S338X mutation. In contrast, both optimized derivatives significantly suppressed phosphorylation in a dose- dependent manner with JM#21 showing the strongest effect (Fig. 5A). Notably, JM#21 was as active as the CXCR4 antagonist AMD3100/Plerixafor. Similar results were observed when phosphorylation of AKT was tested (Fig. 5B). To validate these findings, we also performed kinome profiling of BCWM.1 cells using PamChip technology. This approach allows to simultaneously monitor the enzymatic activity of 144 serine and threonine kinases. In total, 94 kinases were differentially activated between EPI-X4 treated cells and untreated controls. In line with the effects on ERK described above, EPI-X4 suppressed the activation of different members of the MAPK signaling cascade, including p38, ERK1 and ERK2 (Fig. 6).Taken together, these data demonstrate that derivatives of the endogenous CXCR4 inhibitor EPI-X4 have the potential to suppress oncogenic MAP kinase signaling in WM cells expressing wildtype or mutated CXCR4.

**Fig. 5.**
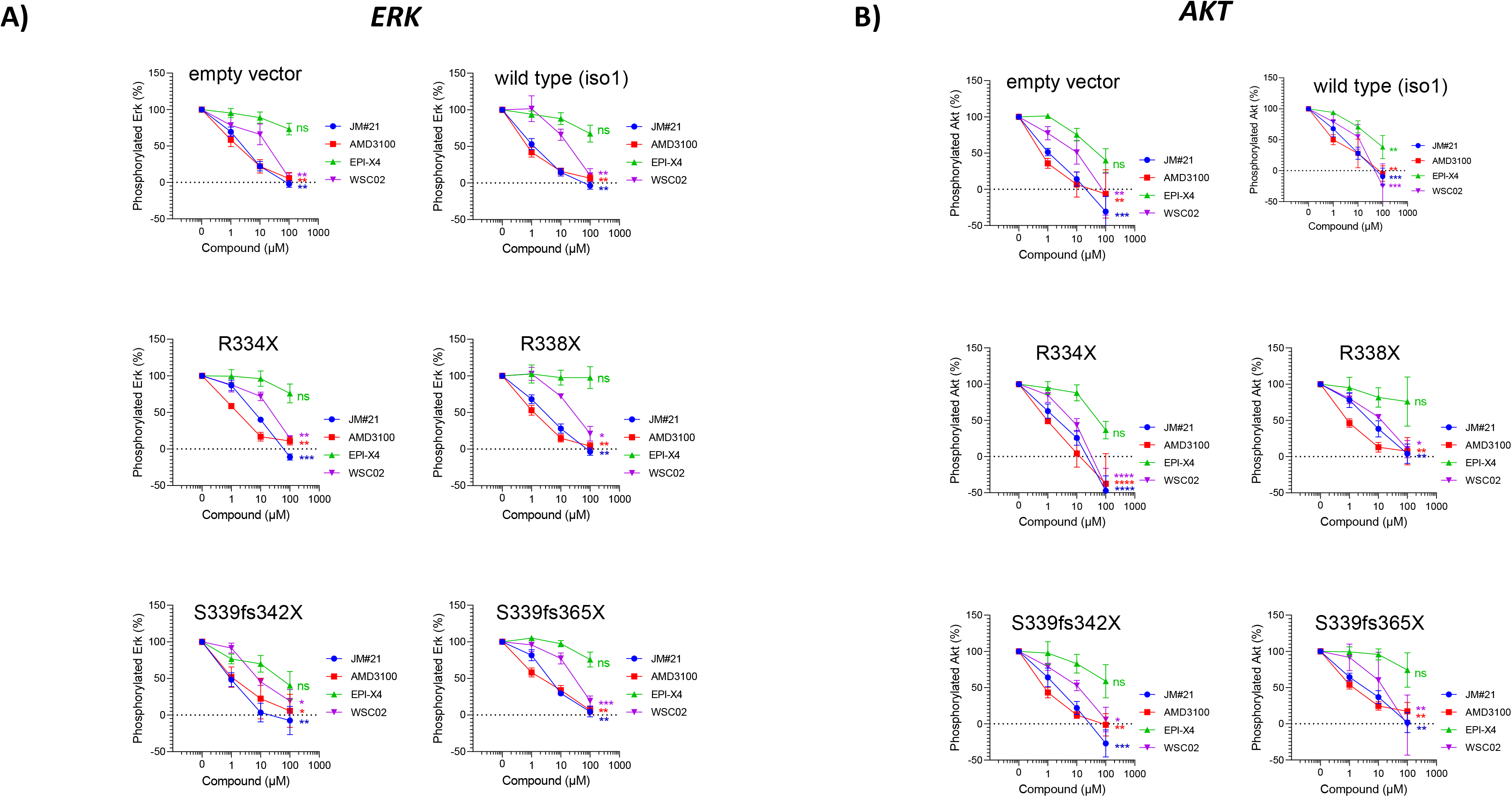
Inhibition of ERK and AKT signaling by EPI-X4 and its optimized derivatives in WM cells. **(A)** ERK and **(B)** AKT phosphorylation in WM cells upon incubation with EPI-X4 and its optimized derivatives. Cells were stimulated with CXCL12 and incubated with the different peptides. Cells were analyzed by flow cytometry with FACS Cytoflex for phosphorylation of Erk1/Erk2 (n=3-5). (* p < 0.1, ** p < 0.01, *** p < 0.001, **** p < 0.0001, ns = not significant)

**Fig. 6.**
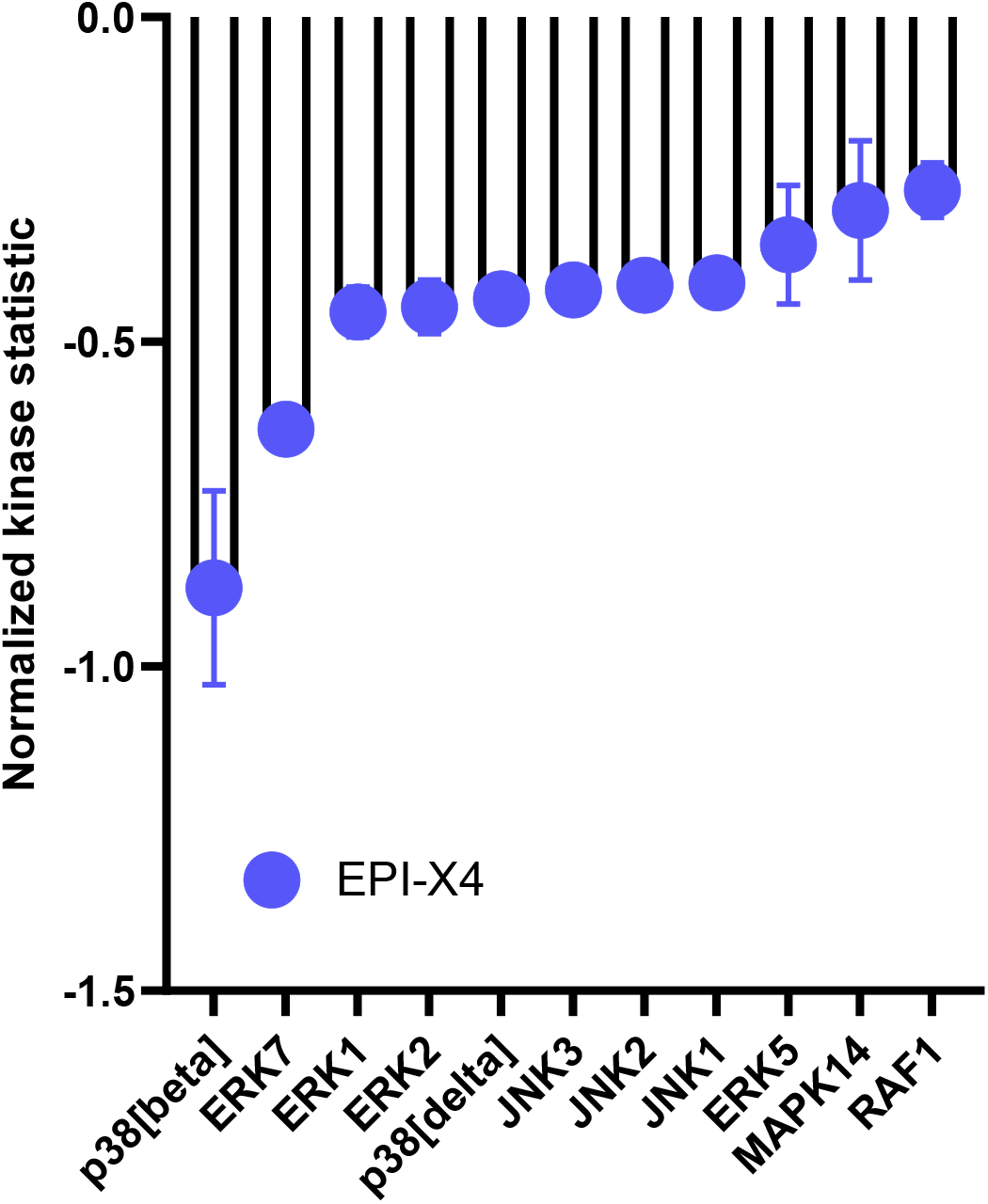
Kinomic changes in MAPK signaling of EPI-X4 treated BCWM.1 cells. Relative kinase activity in EPI-X4 treated vs. untreated BCWM.1 cells. Kinases were ranked using a combined score based on a sensitivity score indicating group difference and specificity score derived for a set of peptides related to the respective kinase. This information is extracted from current experimentally derived or *in silico* databases (BN6). Shown are candidates with a relative kinase activity of −0.25 compared to untreated cells including top candidates related to the MAPK pathway with p38 (delta), p38 (beta) and ERK family members 1, 2 and 5 with a specificity-score of < 1.3.

To further understand the mechanism how EPI-X4 affects gene expression of Waldenström cells, we performed RNA-Seq on the WM cell line BCWM.1 incubated with either EPI-X4 or an inactive control peptide for 24h in serum-free conditions. In total, 4527 genes were differentially expressed, with upregulation of 1920 and downregulation of 2607 genes (Suppl. Table 1). Fold change of gene expression was substantially higher for downregulated genes with up to −1.5 log2 fold differences compared to the inactive peptide control. The top dowregulated gene was *MIF1*, a known ligand of CXCR4 [24]. Furthermore, *IRF7* was among the top 25 downregulated genes This transcription factor is involved in MYD88-associated signaling [25]. *Mir-650* was also among the top 10 downregulated genes and reported to be a prognostic factor in CLL that is associated with tumor invasion and metastases [26]. A similar function has been ascribed to *TSPO*, the third most downregulated gene, additionally known to be highly expressed in lymphoma cells and to induce resistance to apoptosis and H_2_O_2_-induced cytotoxicity in hematopoietic cell lines [27,28]. Strikingly, multiple members of the core subunit of the mitochondrial membrane respiratory chain NADH dehydrogenase (Complex I) such as *NDUFS7*, which was among the top 10 downregulated genes, were downregulated (Supp. Table 1-2, Fig. 7A). Of note, in a first experiment WM cells showed a glycolytic phenotyp and EPI-X4 induced a shift towards aerobic glycolysis as determined by measuring mitochondrial respiration and glycolytic activity using Seahorse Extracellular Flux Analyzer at the early time point of 1, 6 and 24h incubation (Suppl. Fig. 3). When pathways altered by EPI-X4 expression were analyzed by Enrichr, downregulated pathways were enriched for genes involved in Toll receptor, NF-κB, MAPK, AMPK, mTOR and CXCR4/chemokine mediated signaling. Of note, these terms were highly reproducible independent of the libraries used for pathway identification (Fig. 7B, Suppl. Table 2). Upregulated pathways recurrently emerging were FAS, p53, Ubiquitin proteasome and DNA repair/DNA damage pathway (Fig. 7B, Suppl. Table 3).

**Fig. 7.**
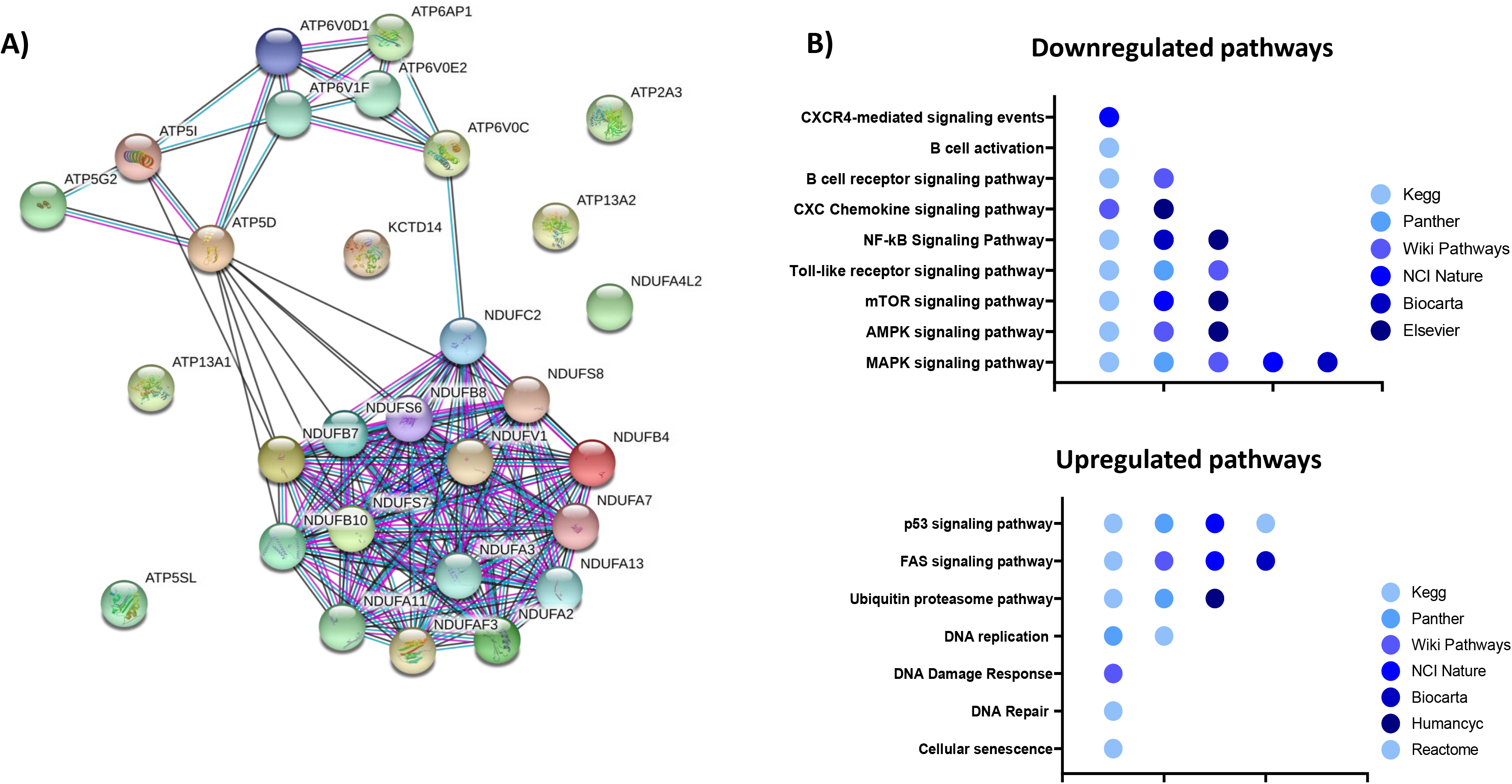
Impact of EPI-X4 on gene expression. **(A)** String analysis of significantly downregulated genes (FDR<0,05) involved in energy metabolism. **(B)** Pathway analysis of differentially expressed genes upon EPI-X4 treatment compared to inactive peptide control (FDR<0,05). Gene set enrichment tools implemented in Enrichr http://amp.pharm.mssm.edu/Enrichr/) were used for the analyses. All gene sets were filtered with an adjusted p-value <0.05.

## Discussion

G protein-coupled receptors (GPCRs) represent the largest family of membrane receptors with an estimated number of 800 members in humans. Several of the GPCRs are known to be critically involved in tumorigenesis. Although GPCRs are targets for approximately 30% of all marketed drugs, only a very limited number of agonists or antagonists acting through these receptors are currently used for cancer therapy [29,30]. CXCR4 has been recognized as a key regulatory mechanism for cancer growth and invasion for a long time and has emerged as a promising target for anti-cancer drugs. In WM, whole genome sequencing lead to the identification of highly recurrent somatic mutations in two genes, namely MYD88 and CXCR4, which has paved the way to a deepened understanding of the signaling cascades driving growth and importantly also therapeutic resistance in WM [31]. The second most recurrent mutation in WM hits the CXCR4 gene: mutations in this gene can be found in up to 40% of the patients. In total more than 40 mutations were described so far [32], all exclusively found in the regulatory cytosolic domain stretching from amino acid position 308 to 352. Almost all mutations described in WM impair CXCR4 desensitization and internalization, thereby prolonging signaling upon binding of the chemokine ligand CXCL12 [14,15]. The main signaling axis that is affected by CXCR4 mutations promote enhanced AKT and subsequent MAPK 1/2 signaling, resulting in sustained survival signals for cancer cells [33]. Patients with CXCR4 mutations are confronted with the downside of showing higher bone marrow involvement, higher IgM levels, symptomatic hyperviscosity and a more aggressive disease at diagnosis with reduced sensitivity towards the BTK – inhibitor ibrutinib. In CXCR4 patients delayed time to response, less deep responses and shorter PFS upon treatment with ibrutinib was observed [32]. Based on all this, it is not surprising that there are major efforts to develop compounds targeting CXCR4 and clinical testing of CXCR4 antagonists have been reported among others in AML and WM [34]. In this context it is, however, surprising that very few is known about naturally occurring mechanisms which are able to directly counteract the evolutionarily highly conserved CXCR4 effects and to dampen or shape CXCR4 signaling in vivo in normal but also in malignant hematopoiesis. Thus, a very important step has been the discovery of EPI-X4, which as a natural CXCR4 antagonist effectively blocks CXCL12-mediated receptor internalization and suppresses the migration and invasion of cancer cells towards a CXCL12 gradient [35]. As we showed there is high expression of CXCR4 at the RNA level in the majority of primary WM samples. However, there is a clear reduced expression in patients with non-mutated MYD88, highlighting the biological difference between MYD88^Mut^ versus MYD88^WT^ patients and suggesting that these patients are not preferentially candidates for CXCR4 targeting. On the other side, these data demonstrate that the vast majority of WM patients could theoretically benefit from CXCR4 targeting. Interestingly, CXCR4 mutated cases showed a significantly increased expression of *HIF1a* and *MAPK1*. It was previously shown that MAPK is required for the transactivation activity of HIF1α and that MAPK signaling facilitates HIF activation through p300/CBP [36]. HIF1α itself was shown to be important for stabilization of CXCR4 expression being recruited to the CXCR4 promoter under hypoxic condition [37]. In summary, this could result in a positive loop, in which CXCR4 mutations would induce increased MAPK signaling, by this supporting HIF1α activity, which itself would lead to stabilization of CXCR4 expression. Based on this and taking the more aggressive clinical course and reduced sensitivity towards BTK inhibition by ibrutinib into account, CXCR4 mutated WM patients would particularly qualify as candidates for a CXCR4 targeting strategy. We could confirm that in particular the most frequent mutation S338X increases growth of WM cells and this independently of the microenvironment. EPI-X4 was able to bind to WM cells and importantly impaired significantly clonogenic growth and migration in vitro. One underlying mechanism for this is impairment of MAPK signaling, known to be vital for WM cell growth. Beside this, we could demonstrate that binding of EPI-X4 changed the transcription of genes involved in energy metabolism, becoming particularly evident by reduction of expression of multiple members of the core subunit of the mitochondrial membrane respiratory chain NADH dehydrogenase (Complex I) such as *NDUFS7*. In addition, we also observed reduction of *NAMPT* transcription (data not shown), all indicating that the naturally occurring CXCR4 antagonist is able to dampen the energy metabolism of WM cells. However, at least at an early time point we did not see a major drop in glycolysis but a shift towards anaerobic glycolysis testing mitochondrial respiration in these cells. Furthermore, when pathways altered by EPI-X4 expression were analyzed by Enrichr, downregulated pathways were also enriched for genes involved in pathways related to metabolism such as glycolysis, pentose phosphate pathway and OXPHOS (data not shown). Data on metabolism in WM are very limited and just recently it was reported that WM cells are enriched for glutathione metabolism based on unsorted WM bulk derived from peripheral blood or BM compared to normal B-cells [38]. Data to which extent CXCR4 antagonists impact the metabolomic fingerprint in WM are not existing. In light of this, nicotinamide metabolism has emerged as one major topic as well as dependencies of cancer stem cells on amino acid metabolism. NAMPT overexpression has been described in cancer cells and pharmacological NAMPT targeting has shown anti-tumor effects in AML and WM [39,40]. Just recently, resistance to Venetoclax/Azacytidine in AML has been linked to increased nicotinamide metabolism at the level of AML LSCs [41]. One of the key questions is whether the discovery of EPI-X4 can fuel the development of novel CXCR4 antagonists for the treatment of WM and other CXCR4 dependent malignancies. In contrast to AMD3100, the only clinically approved CXCR4 antagonist, EPI-X4 also reduces basal CXCR4 signaling in the absence of CXCL12 and does not interact with CXCR7, whereas AMD3100 acts as an allosteric agonist of this receptor. Moreover, EPI-X4 and improved derivatives thereof do not exert mitochondrial cytotoxicity, in contrast to AMD3100 [19]. However, we observed only moderate effects of EPI-X4 in some experiments, and the half- life of the peptide is short with about 17 min in serum- containing medium. Thus, generation of optimized derivatives of EPI-X4 will be crucial for opening the door to clinical application. We performed systematic, quantitative structure activity relationship (QSAR) studies to further improve the anti-CXCR4 activity of EPI-X4 and have solved NMR structures of EPI-X4, as well as first (WSC02) and second (JM#21) generation derivatives, and have used this information for computational modeling to elucidate their exact binding mode with CXCR4, and to predict peptides with further improved activity [23]. In iterative processes, we have now synthesized and tested more than 150 EPI-X4 derivatives (data not shown), enabling the identification of peptides with activities in the low nanomolar range. Indeed, WSC02 and JM#21 showed an enhanced activity in several assays compared to EPI-X4 in WM, underlining the potential of optimized derivatives to serve as novel anti-WM compounds in pre-clinical and later on clinical testing. Taken together, our data point to a yet unknown naturally occurring mechanism to dampen CXCR4 activity in WM and the potential to translate these insights into the biology of WM into the development of promising novel therapeutic compounds in this indolent lymphoma subtype.

## Materials and Methods

### Quantification of CXCR4 surface levels by flow cytometry

All cell line experiments were conducted under serum-free conditions (RPMI plus 1% penicillin/streptomycin). Cells were incubated with different CXCR4 antagonists or an inactive control peptide at different concentrations as indicated for 30 min at 4 °C. A control sample was incubated with PBS only to determine basal receptor levels at the cell surface. After incubation, cells were stained with anti-CXCR4 antibodies (APC-labeled 12G5 antibody, 555976 or PE-labeled 1D9 antibody, 551510 from BD Biosciences).

### Stable transduction of BCWM.1 and MCWL-1 cells

Retroviral particles were generated by cotransfecting Lenti-X™ 293T cells with the lentiviral vector pCDHMSCV-EF1-GFP-T2A-PURO expressing CXCR4 iso1 WT (isoform 1 or b, NCBI Reference Sequence: NP_003458.1,UniProtKB/Swiss-Prot: P61073-1) or CXCR4 mutants, the packaging plasmid pSPAX2, and VSV-G encoding pMD2.G, using TransIT®-LT1. Cells were sorted twice on a BD FACSAria™ III for the fluorescence marker APC (CXCR4) and GFP expressed from pCDH-MSCV-EF1-GFP-T2A-PURO. As WM cell lines BCWM.1 and MCWL-1 cells were used (both MYD88 mutated/CXCR4 wildtype).

### CFC assay

Colony forming cell unit assay was performed as previously described [42]. 4000 cells were plated per dish for WM cell lines (Methocult H4330 StemCell Technologies). Colonies were scored on day 7 after plating.

### Migration

BCWM.1 cells constitutively expressing CXCR4 iso1 WT, CXCR4 S338x or the empty vector control were seeded together with EPI-X4, WSC02, JM#21 or an inactive peptide control in the upper well of a transwell. 10 nM CXCL12 was supplemented to the lower chamber. After 2h incubation at 37°C, 5% CO_2_, cells that had migrated to the lower compartment were analyzed using the CellTiter-Glo Luminescent Cell Viability Assay according to the manufacturer’s instructions. Percentage of specific migration was calculated using the following formula: % migration= ((specific migration-unspecific migration)/total cells) *100.

### Total RNA extraction and RNA sequencing

BCWM.1 cells were cultured in medium without FBS containing 200 μM of either the inactive peptide or EPI-X4 as well as 100 ng/ml CXCL12. After 24 h, cells were collected, and RNA was isolated using the RNA isolation Kit Direct-zol RNA MiniPrep according to the manufacturer’s protocol. After quality check, the RNA was prepped using the TruSeq RNA sample Kit from Illumina, and samples were run on the Illumina HiSeq 2000 sequencer. Afterwards, alignment were performed, and gene expression was analyzed by basepair.

### Kinome profiling

Cells were lysed using M-PER™ Mammalian Protein Extraction Reagent (78503, Thermo Scientific) supplemented with Halt™ Protease Inhibitor Cocktail (87785, Thermo Scientific) and Halt™ Phosphatase Inhibitor Cocktail (78420, Thermo Scientific). Protein concentration was determined using Qubit. Kinase activity in cleared lysates was assessed using serine-/threonine-kinases (STK) assay (PamGene International BV). Reactions were carried out in the presence of 400 μM ATP in combination with the STK reagent kit and 2 μg total protein. Samples were applied to porous aluminum oxide arrays spotted with 144 immobilized 13-amino acid peptide substrates containing STK phosphosites. The pumping of samples through the arrays and imaging of FITC-coupled antibodies detecting phosphorylated peptides on the arrays at different exposure times via a CCD camera were carried out using the PamStation12 instrument operated with the Evolve software (PamGene International BV). Subsequent image quantification and internal quality-control was performed using BioNavigator 6 ® software (BN6, PamGene International BV) with resulting signal-intensities per peptide being log2-transformed. A functional scoring tool Upstream Kinase Analysis (BN6) is used to generate a putative list of kinases responsible for phosphorylating the phosphosites on the PamChip.

### Metabolic Assessment

After pretreatment with the compound in the indicated concentrations and timepoints, BCWM.1 cells were plated on 96 well cell culture (XFe96, Agilent Technologies, Santa Clara, USA) which had been precoated with CellTak (Thermo Fisher Scientific, Hampton, USA) in bicarbonate-free RPMI medium containing 1 mM HEPES, 10 mM glucose, 1 mM pyruvate, and 2 mM glutamine. Oxygen consumption and extracellular acidification rates (OCR and ECAR) were measured simultaneously using a Seahorse XFe96 Flux Analyzer (Agilent Technologies). Uncoupled (proton leak) respiration was profiled by injecting 1.5 μM oligomycin (inhibiting the ATP synthase). Non-mitochondrial respiration was determined by injecting 0.5 μM antimycin A and 0.5 μM rotenone (inhibiting electron flux through complex I and III). OCR and ECAR were determined by machine algorithms and plotted against time. Data were normalized to cell content by staining with Janus Green [43]. ATP production rates from oxidative phosphorylation were calculated assuming a phosphate/oxygen (P/O) ratio of 2.75. ECAR were converted into proton efflux rates (PER) which were used for calculating glycolytic ATP production rates in a 1:1 ratio.

### Mouse experiments

All mouse experiments were conducted according to the national animal welfare law (Tierschutzgesetz) and were approved by the Regierungspräsidium Tübingen, Germany. For all experiments, NOD.Cg-Prkdc<scid>Il2rgytm1Wjl>/SzJ (NSG) strains were used. The NSG mice were irradiated sublethally (325 cGy) and injected with MCWL-1 cells incubated with 2 mM peptides for 30 min under serum-free condition.

### Statistics

All statistical analyses were performed using the GraphPad Prism 7 software using Ordinary one-way ANOVA.

## Acknowledgments

We want to thank Olivier Bernard (INSERM U1170 and Gustave Roussy, Villejuif, France) for providing RNA-Seq data of WM patients. The authors would like to thank all members of the Core Facility FACS and the animal facility of the University Ulm for breeding and maintenance of the animals. The work was supported by grants from the DFG (SFB 1279 project B01 to C.B, and project A04 to M.H. and J.M.; SPP 1923 to D.S.) and by a grant of the International Waldenström Foundation (IWMF) to J.M., D.S. and C.B. M.H. is member of the International Graduate School in Molecular Medicine Ulm.

## Author Contributions

L.M.K., M.H., M.G, Z.H. and V.P.S.R. performed research and analysed the data. J.M., D.S. and Z.H. contributed to data interpretation. L.M.K. and C.B. wrote the manuscript, C.B. designed the project.

## Supplemental Information

**Supplemental Tables:**

**Supplemental Table 1**: Differentially expressed genes between BCWM.1 cells treated with EPI-X4 for 24 h (200 μM) versus inactive peptide. Genes with a FDR < 0,05 and p< 0.05 are listed.

**Supplemental Table 2:** Pathway analysis of differentially expressed and downregulated genes between BCWM.1 cells treated with EPI-X4 for 24 h (200 μM) versus inactive peptide using Enrichr. Genes which are involved in the pathways depicted in Fig.9 B are marked in yellow (adjusted p-value < 0.05).

**Supplemental Table 3:** Pathway analysis of differentially expressed and upregulated genes between BCWM.1 cells treated with EPI-X4 for 24 h (200 μM) versus inactive peptide using Enrichr. Genes which are involved in the pathways depicted in Fig.9 B are marked in yellow (adjusted p-value < 0.05).

**Supplemental Figures:**

**Supplemental Figure 1:**
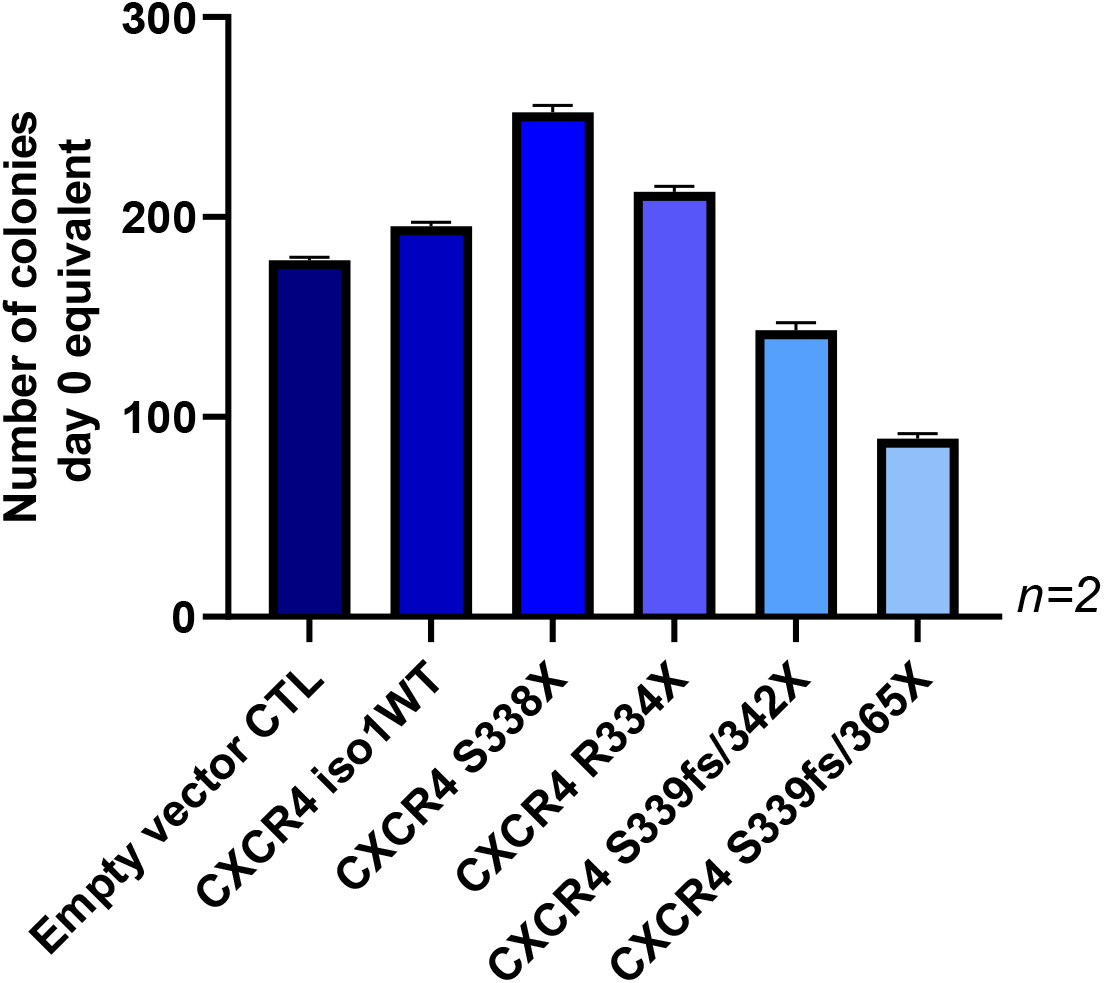
Colony formation ability of BCWM.1 cells transduced with CXCR4 S338X, R334X, S339fs/342X and S339fs/365X as well as the iso1 WT control. Mean values (± SEM) of two independent experiments performed in duplicates are shown.

**Supplemental Figure 2:**
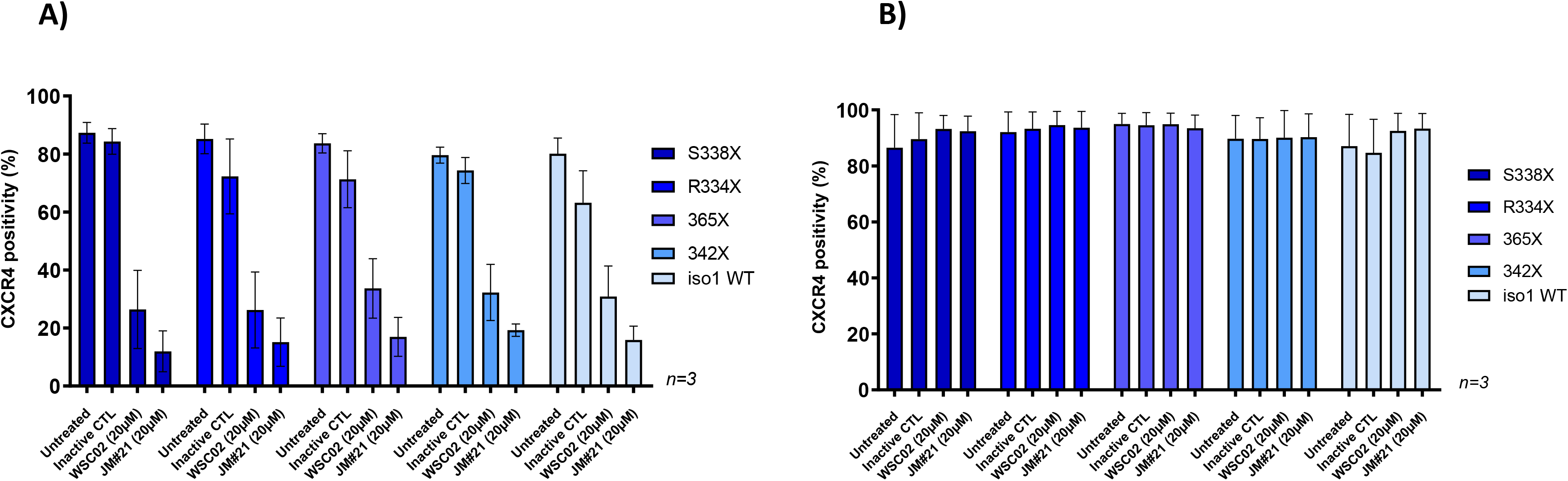
Successful blockage of all CXCR4 mutants in BCWM.1 cells by low concentrations of optimized derivatives of EPI-X4. BCWM.1 cells were preincubated with 20 μM WSC02, JM#21 or inactive control peptide and then stained with the anti-CXCR4 antibody clones 12G5 (A) or 1D9 (B). While both EPI-X4 derivatives block binding of the anti-CXCR4 clone 12G5 to all CXCR4 variants tested, they do not affect binding of the 1D9 clone. Mean values (± SEM) of one exemplary experiment performed in triplicates are shown.

**Supplemental Figure 3:**
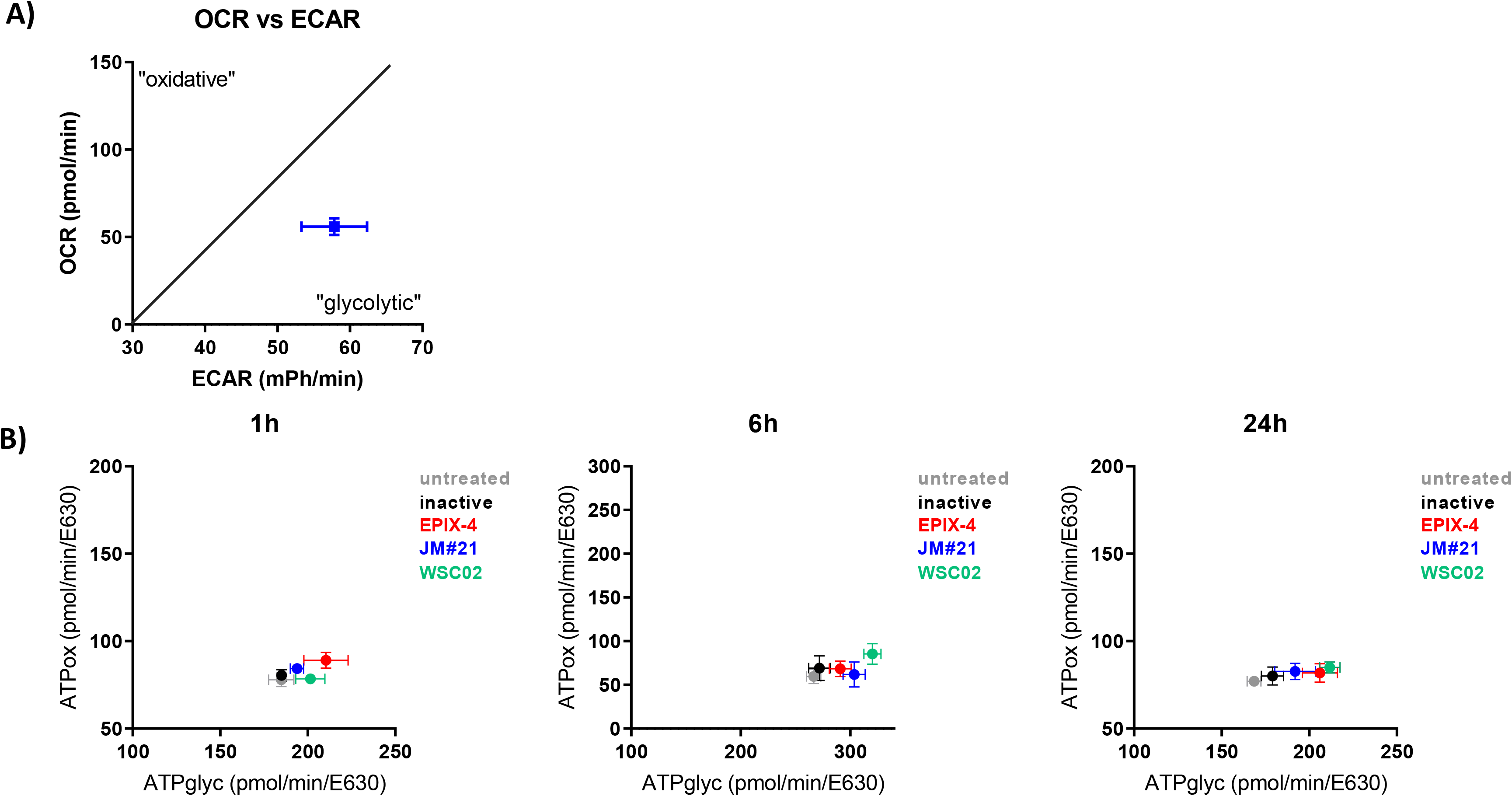
Metabolic assessment using Seahorse XF. **(A)** BCWM.1 cells were cultivated in normal growth medium (RPMI 10% FBS 1% Penicillin/Streptomycin) and mitochondrial function was assed measuring oxygen consumption rate (OCR) and extracellular acidification rate (ECAR) **(B)** BCWM.1 cells were cultivated for 1,6 and 24 h under serum-free conditions in the presence or absence of the indicated CXCR4 inhibitors (inactive peptide: 10 μM, EPI-X4: 200μM, WSC02: 10 μM, JM#21: 10μM). Cells were evaluated in terms of their mitochondrial function measuring basal respiration and oligomycin-induced changes in oxygen consumption rate (OCR) and extracellular acidification rate (ECAR), thereby calculating ATP production rates.

